# nf-core/marsseq: systematic pre-processing pipeline for MARS-seq experiments

**DOI:** 10.1101/2023.06.28.546862

**Authors:** Martin Proks, Jose Alejandro Romero Herrera, Jakub Sedzinski, Joshua M. Brickman

## Abstract

**Motivation:** As a result of advancing single sequencing technology (scRNA-seq), it has become possible to study gene regulatory mechanism(s) and their influence on evolving cell states in time at the level of individual cells. Since 2009, numerous scRNA-seq protocols have been developed, each with its own advantages, disadvantages and library preparation complexities (Ziegenhain et al. 2017). However, the interpretation of data arising from these techniques often shares similar limitations, such as the lack of a standardized pre-processing workflow and consistent data reproducibility. Here we focus on the standardization of the plate based Massively Parallel RNA Single cell Sequencing (MARS-seq, Jaitin et al. 2014) pre-processing pipeline as described in MARS-seq2.0 (Keren-Shaul et al. 2019), which was developed at the Weizmann Institute of Science.

**Results:** To overcome the limitations mentioned above, we have taken the original MARS-seq2.0 pipeline and revised it to enable implementation using the nf-core framework (Ewels et al. 2020). By doing so, we have simplified pipeline execution enabling streamlined application, with increased transparency and scalability. Additionally, we have further improved the pipeline by implementing a custom workflow for RNA velocity estimation.

**Availability and implementation:** The pipeline is part of the nf-core bioinformatics community and is freely available at https://github.com/nf-core/marsseq with data analysis at https://github.com/brickmanlab/proks-et-al-2023.

## 1. Introduction

Over the past decade, several new protocols for single-cell RNA sequencing (scRNA-seq) have been developed focusing on the collection of transcriptomic data via different approaches, each with specific advantages relevant distinct experimental questions. With time, an increased number of novel approaches have been developed to study mRNA abundance at a single cell resolution. Popular methods include cell isolation using microfluidic devices, plate-based techniques, and nanopore-based strategies (Mereu et al. 2020). Each technique requires an additional pipeline that: i) pre-processes raw reads, ii) aligns them to a reference genome, iii) performs demultiplexing to match individual transcript to specific cells, and iv) reports on the number of expressed genes per individual single cell in a tabular format known as count matrix.

Microfluidic methods that use unique molecular identifiers (UMI) tag systems are popular and can be processed by open-source tools, such as, dropEst (for inDrop, Drop-seq) (Petukhov et al. 2018) kallisto bustool (for UMI generic) (Melsted et al. 2021) or umis (for UMI generic) (Svensson et al. 2017). Commercial platforms like 10X Chromium offer an out-of-the-box tool called cellranger (Zheng et al. 2017). Plate-based methods like full-length SMART-seq2/3 or MARS-seq can be processed using zUMIs (Parekh et al. 2018) or StarSolo (Kaminow, Yunusov and Dobin 2021). To fully automate sequencing read pre-processing, these tools can be wrapped in a pipeline using workflow managers like snakemake (Mölder et al. 2021) or Nextflow (Di Tommaso et al. 2017), to take full advantage of high-performance computing (HPC) resources.

In this paper we focus on the plate-based MARS-seq protocol, that incorporates fluorescence-activated cell sorting (FACS) to investigate rare populations of cells. This approach reduces the occurrence of cell doublets and neighbouring cell contamination, while keeping experimental costs low. We use two datasets with different contours; a large dataset which spanned 33,700 cells across 153 embryos that produced new insights into mouse gastrulation and a smaller dataset that focused on the differentiation of the endoderm lineage, identifying a rare, but pivotal transition state in lineage specification.

Unfortunately, setting up and executing the original pipeline for MARS-seq using MARS-seq2.0 proved to be a challenging task, as the pipeline required customizing additional configurations for aligners and manually generating metadata files for each experiment. Despite thorough documentation, the setup and execution of the pipeline had its own difficulties. To avoid these problems and make this technology more accessible we streamlined the execution for non-computational researchers and enhanced its portability.

## 2. Materials and Methods

The nf-core/marseq pipeline is straightforward to execute and involves two main steps. First, the building of the necessary reference indexes for a designated genome (in this case, either human or mouse). Second, the pipeline aligns the raw reads and generates a count matrix that is then utilized for further downstream analysis.

Our pipeline was developed utilizing the nf-core/tools template (Ewels et al. 2020), a community-curated python package adhering to best-practice and employing Nextflow as a underlying workflow manager. Nextflow’s new domain-specific language (DSL2) syntax provides a modular structure to the code, making it easy to extend the pipeline in future releases. Taking advantage of Nextflow, the pipeline can be executed on local computer, High Performance Computing (HPC) and cloud providers (such as Amazon and Google). Tracking of the pipeline can be done online using the open-source external service Nextflow Tower (Seqera Labs 2019). Container technologies such as Docker, Podman, or Singularity are also supported to ensure future reproducibility of the pipeline and pre-processed data. Lastly, nf-core/marsseq implements extra steps that enable the estimation of RNA velocity, which was previously not possible with existing MARS-seq2.0 pipelines. A summary of all available workflows can be found in **Figure 1**.

**Figure 1:**
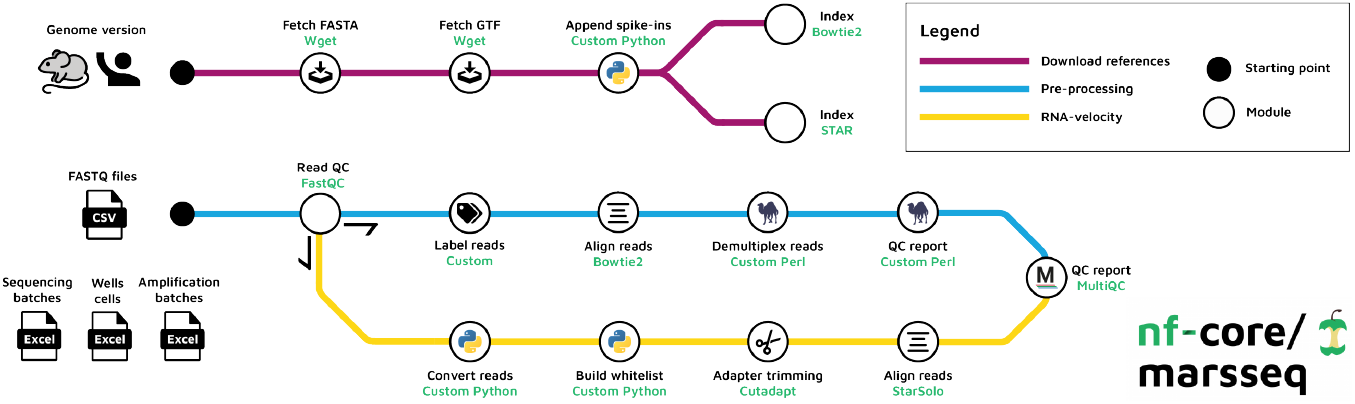
nf-core/marseq supports both mouse and human genomes. It consists of three workflows: i) reference building, ii) construction of count matrix, and iii) RNA velocity estimation. At the end of each run, all quality control statistics are summarized with MultiQC.

### 2.1. Reference building

The original MARS-seq2.0 pipeline provides only one custom mouse (mm9) reference for download. The custom reference is required due to the use of RNA Controls Consortium (ERCC) spikes as controls for accurate gene expression measurements (Pine et al. 2016). The incorporation of these spikes requires conversion of ERCC sequences (n=92) to FASTA format and manually concatenating them to the genome reference, as specified in the MARS-seq2.0 documentation (Tanaylab 2019). In our pipeline we have automate this process, so that we can specify the organism, and then download a reference genome (FASTA), and annotation (GTF) from the GENCODE database. The pipeline then appends the ERCC spike-in sequences and builds reference indexes for required aligners. Bowtie2 (Langmead and Salzberg 2012) is set as the default aligner, following the original MARS-seq pipeline. The workflow is summarised in the upper panel of **Figure 1**. Reference building can be executed using a simple one-line command:

~~~
nextflow run nf-core/marsseq -profile test,YOURFILE --build_references --genome mm10 --velocity
~~~

### 2.2. Pipeline execution

Running the nf-core/marsseq pipeline requires the use of custom references from the previous step, as well as a design file that contains information about the specific experiment. Each MARS-seq experiment is comprised of paired end raw (FASTQ) reads and one or multiple sequencing batches (SB) that are described in the seq_batch.txt file. Each SB contains multiple amplification batches (AP, amp_batch.txt), and the wells contained within that batch (WS, described in wells_cells.txt). The original pipeline requires these files in tabular separated format (tsv). However, based on our experience, it is much easier to create and update these files using Microsoft Excel (xlsx). Therefore, we have developed a set of custom Python scripts that can convert Excel files into the required tabular format and validate the resulting files, checking for any mistakes. If a mistake is detected, the pipeline stops and notifies the user with an appropriate error message. These additional validation steps were not included in the original MARS-seq2.0 workflow and therefore produced inconsistent error messages without terminating pipeline execution. To maintain consistency with previous pipeline, we have kept the metadata changes to a minimum. Our new pipeline documentation includes a prefilled metadata template to facilitate its use.

Assuming that all input files are correct, the pipeline first performs parallel quality control on raw reads using FastQC (Andrews et al. 2012) and assigns barcode labels to the sequence reads. The FASTQ files are then divided into subsets of 4,000,000 reads per file to avoid bowtie’s memory restrictions during alignment. Poor-quality reads are discarded while the rest are trimmed (removal of adapters). By default, alignment is performed using the bowtie2 aligner. The aligned reads are subsequently demultiplexed based on labelled barcodes, followed by the generation of a count matrix. Finally, the quality control (QC) reports are generated and summarized using MultiQC (Ewels et al. 2016). To convert the original pre-processing Perl scripts into Nextflow modules, we had to make some additional changes. Unless specified otherwise, all output is stored in the *results* folder. The nf-core/marsseq pipeline can be executed as follows:

~~~
nextflow run nf-core/marsseq -profile test,YOURFILE --input samplesheet.csv
~~~

### 2.3. RNA Velocity estimation

RNA velocity is a powerful method for predicting cellular dynamics and future cell states during differentiation by modelling the relationship between the observed number of spliced and unspliced RNAs (Bergen et al. 2020). This process, however, requires a splice aware aligner such as kallisto, Alevin-fry (He et al. 2022) or StarSolo. Another option is to use already aligned bam files and process them with the Python package velocyto (La Manno et al. 2018). Unfortunately, none of these tools support MARS-seq double barcoding. To overcome this limitation, we have developed a set of Python scripts that convert MARS-seq reads to 10X format.

MARS-seq is a paired-end method where read 1 (R1) consists of a left adapter (LA, 3 bs), a pool barcode (PB, 4 bs) and cDNA (68 bs). Read 2 (R2) contains a cell barcode (CB, 7 bs) and a UMI (8 bs). All barcodes are stored in the header of each individual read. On the contrary, 10X R1 consists of a CB and a UMI, while the cDNA is stored in R2. To mimic the 10X format, we merge PB, CB and UMI to generate R1 and move the trimmed (adapter free) cDNA to R2, as illustrated in **Figure 2**.

**Figure 2:**
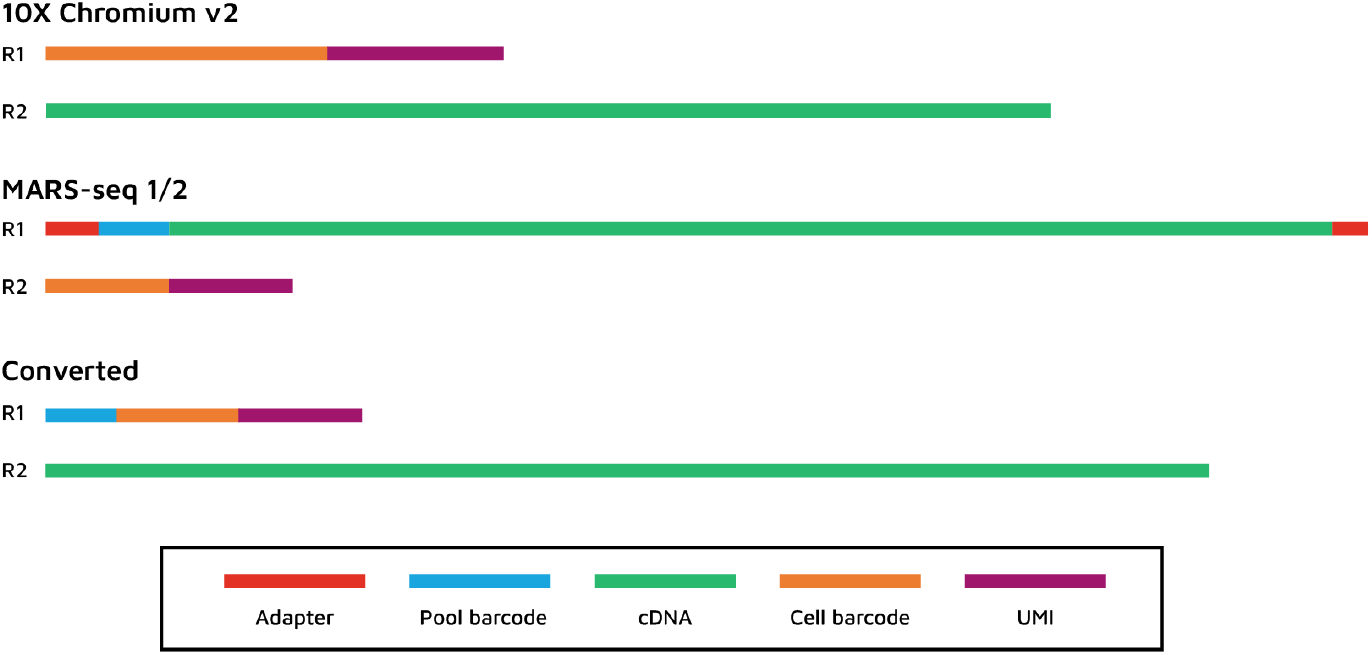
10X) R1: CB (16bp) + UMI (10bp); R2: cDNA (57bp). MARS-seq) R1: LA (3bp) + PB (4bp) + cDNA (66bp) + RA (2bp); R2: CB (7bp) + UMI (8bp). Converted) R1: PB (4bp) + CB (7bp) + UMI (8bp); R2: cDNA (66bp). Legend: CB (cell barcode), UMI (Unique Molecular Identifier), LA (left adapter), PB (pool barcode), RA (right barcode).

Based on previous work that attempted to benchmark different tools (Soneson et al. 2021, Du et al. 2020), we have decided to use StarSolo for estimating unspliced reads. To demultiplex the reads, the pipeline generates a file containing all valid cell barcodes containing concatenated PB with CB (11 bp) called whitelist.txt, which is required by StarSolo. We have also adjusted the default parameters to comply with the updated barcoding scheme as follows: *--soloType CB_UMI_Simple --soloCBstart 1 --soloCBlen 11 --soloUMIstart 12 --soloUMIlen 8 --soloFeatures Gene GeneFull SJ Velocyto*. To enable the estimation of spliced, unspliced and ambiguous transcripts, one simply need to append the below *velocity* flag:

~~~
nextflow run nf-core/marsseq -profile test,YOURFILE --input design.csv --velocity
~~~

## 3. Results

To validate the reliability of our pipeline, we tested it on two small datasets (SB26 and SB28) provided in the MARS-seq2.0 documentation, alongside the original MARS-seq2.0 pipeline. Both pipelines used mouse (mm10) as the reference genome for mapping. We found that the read counts and number of genes per individual cell were highly similar between the two pipelines (ρ = 0.999) (**Figure 3A**).

**Figure 3:**
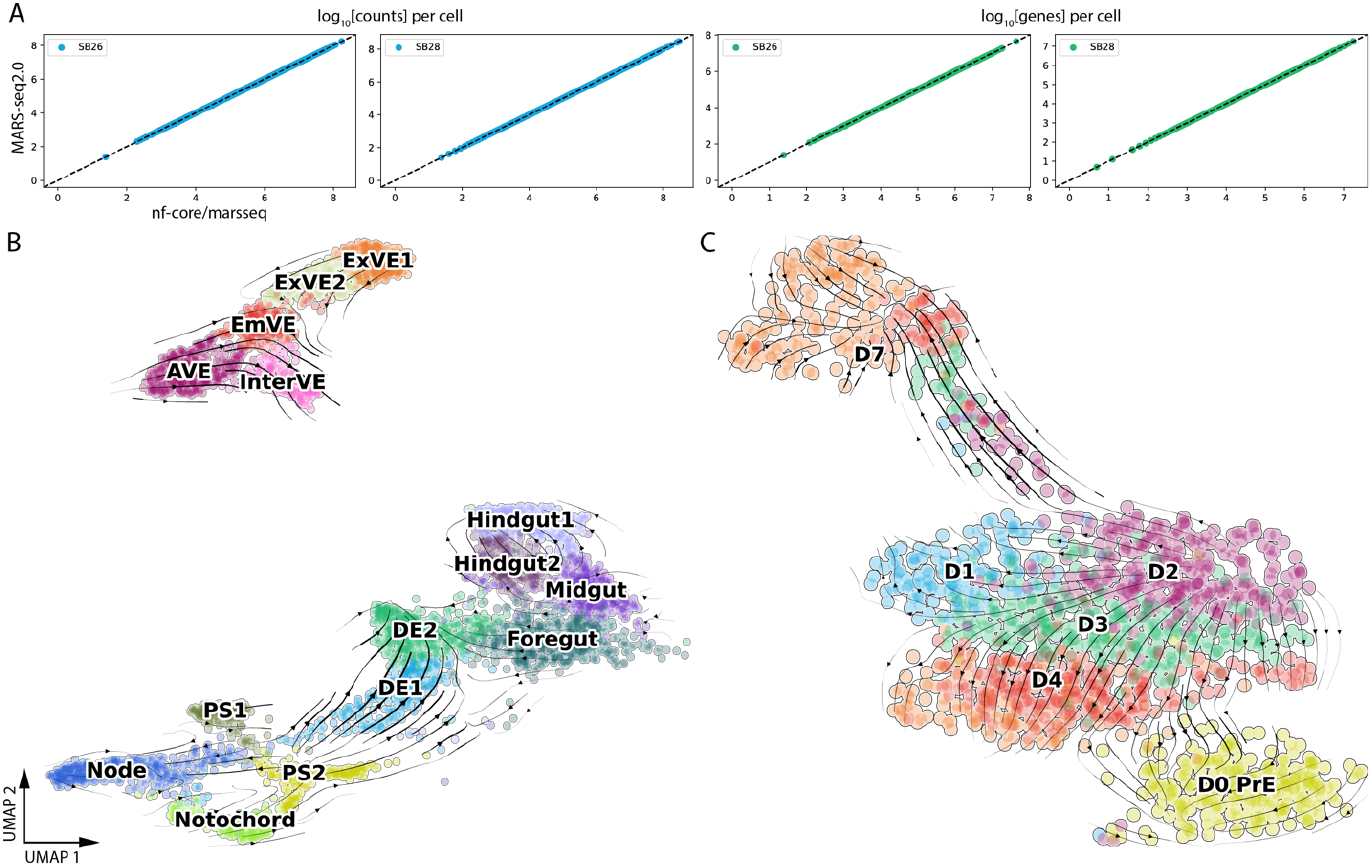
**A)** Scatter plot comparing number of counts per cell (left blue panels) and number of genes detected per cell (right green panels) between the original MARSseq2.0 and nf-core/marseq pipelines. The dotted line represents a perfect fit (Pearson correlation of 1). **B**) Visualization of RNA velocity computed from (Rothová et al. 2022). **C)** Re-analysis and visualization of RNA velocity estimation of *in vitro* PrE differentiation from (Perera et al. 2022).

Next, we visually validated RNA velocity inference using two datasets; the first example was published in (Rothová et al. 2022) and the second involved the use of published data (Perera et al. 2022) to generate new inferences. In published work, we investigated in vivo endoderm differentiation using a FOXA2^Venus^ reporter to purify endoderm from mouse embryos. We had identified an intermediate cell population that provided a bridge that allowed the conversion of previously thought to be extra-embryonic visceral endoderm, into embryonic endodermal organs. Using trajectory inference (Partition-Based Graph Abstraction, PAGA) and RNA velocity, we were able to describe the route taken by visceral endoderm, across this bridge and onwards. To determine whether this route could be recapitulated in vitro, we differentiated naïve extra-embryonic endoderm stem cells (Anderson et al. 2017), in three dimensional cultures to produce gut spheres, which were characterized by scRNA-seq using MARS-seq. As with the in vivo data, we used trajectory inference, to determine the route of endoderm differentiation in vitro.

At the time of writing (Rothová et al. 2022), the nf-core/marsseq was still in development. We therefore published the RNA velocity workflow as a bash script which was later incorporated into our new pipeline as a module. In short, the original FASTQ files were re-aligned to mm9 using STARSolo v2.7.9a to infer spliced and unspliced reads. Count matrices were merged with the already analysed dataset and further processed using *scVelo* (Bergen et al. 2020). We projected the estimated velocities onto the original UMAP visualization, as shown in **Figure 3B**.

In (Perera et al. 2022) the link between lineage priming, the cell cycle and lineage specification is explored by a combination of imaging and scRNA-seq. We found that during primitive endoderm (PrE) differentiation, the G1 phase of the cell cycle is increased, despite an enhanced rate of cell division. This increase in G1 was observed both in vivo and in vitro for PrE differentiation. The position of cell cycle phases relative to lineage choice was determined by live imaging or by gene expression analysis that utilized a double fluorescent reporter cell line, containing both an epiblast/embryonic stem cell (ESC) specific Sox2-GFP and PrE-specific Hhex-mCherry. ESCs were differentiated in vitro for 7 days to PrE using the protocol described in (Anderson et al. 2017) and sorted based on reporter expression followed by sequencing at different stages of differentiation using MARS-seq. We pre-processed the data with the MARS-seq2.0 pipeline using mm9 genome as reference. We identified groups of cells that either differentiated towards the PrE fate or remained in an epiblast like state (NEDiff) fate. As the original dataset contained equal proportions of each sorted population regardless of its representation in the culture at that time point, the distribution of NEDIFF and PrE cells sequenced did not reflect the increasing proportion of these cultures that were becoming PrE.

To address this issue, we pre-processed the original FASTQ files with nf-core/marsseq with an RNA workflow enabled using mm10 genome. New raw count matrices were merged and processed using *scanpy* (Wolf, Angerer and Theis 2018), while RNA velocity analysis was carried out using *scVelo*, projecting velocities on newly recomputed UMAP. Our new analysis supports time lapse imaging and marker analysis in (Perera et al. 2022), demonstrating a clear bifurcation with two trajectories (**Figure 3C**).

## 4. Discussion & Conclusion

In this paper, we developed a robust and reproducible pre-processing Nextflow pipeline for MARS-seq experiments using nf-core tools. To achieve this, we took the original MARS-seq2.0 pipeline and broke it down into individual pre-processing steps, wrapping them into modules using Nextflow’s DSL2 syntax. Our pipeline consists of two main workflows: reference building and execution. We simplified the required input files as well as integrating a set of checkpoints for their validation, making the pipeline sufficiently simple to run such that it is accessible to molecular biologists without extensive computational experience. By using Nextflow as workflow manager, the pipeline can be executed both on both cloud as well as in-house HPC systems. Additionally, it supports standard containerized solutions that safeguard reproducibly. Lastly, execution can be tracked online by integrating it with Nextflow Tower. At the time of submission of this work we identified a tool that was designed to interconvert different sequencing formats (Battenberg et al. 2022); however, although this appears a robust tool for many applications, we were unable to use it to convert MARS-seq2.0 to 10X format.

Additionally, we have developed a workflow for RNA velocity inference using MARS-seq data. This pipeline has been applied to several datasets in our lab, where it has successfully identified novel differentiation trajectories, both in vivo (Rothová et al. 2022) and in vitro (Perera et al. 2022) (**Figure 3**). This is achieved by converting MARS-seq reads into 10X v2 format and customizing the parameters for the STARSolo aligner as a means to estimate unspliced read counts. This approach has produced important insight into novel routes of endoderm differentiation.

Currently, our pipeline supports the original workflow using the Botwie2 aligner and can generate an additional count table using the RNA velocity workflow. However, based on our experience and findings from (Du et al. 2020), we recommend using the STARSolo aligner for speed. However, the biggest bottleneck of the pipeline is the conversion to 10X format, which is limited by the I/O speed of hard drives.

## Acknowledgments

We thank Eyal David for consulting on execution of the original pipeline. We would also like to thank Maxime U. Garcia for guidance on building custom nf-core modules. This work was supported by the Novo Nordisk Foundation (NNF17OC0028218), the Lundbeck Foundation (R370-2021-617, R198-2015-412 and R286-2018-1534), and the Danish National Research Foundation (DNRF116). The Novo Nordisk Foundation Center for Stem Cell Medicine is supported by Novo Nordisk Foundation (grant number NNF21CC0073729 and previously NNF17CC0027852).

## Contributions

M.P drafted the manuscript and developed the pipeline with assistance of J.A.R.H. J.M.B and J.S wrote the manuscript and supervised the project.

